# iScore: an MPI supported software for ranking protein-protein docking models based on a random walk graph kernel and support vector machines

**DOI:** 10.1101/788166

**Authors:** Nicolas Renaud, Yong Jung, Vasant Honavar, Cunliang Geng, Alexandre M.J.J. Bonvin, Li C. Xue

## Abstract

Computational docking is a promising tool to model three-dimensional (3D) structures of protein-protein complexes, which provides fundamental insights of protein functions in the cellular life. Singling out near-native models from the huge pool of generated docking models (referred to as the scoring problem) remains as a major challenge in computational docking. We recently published iScore, a novel graph kernel based scoring function. iScore ranks docking models based on their interface graph similarities to the training interface graph set. iScore uses a support vector machine approach with random-walk graph kernels to classify and rank protein-protein interfaces.

Here, we present the software for iScore. The software provides executable scripts that fully automatize the computational workflow. In addition, the creation and analysis of the interface graph can be distributed across different processes using Message Passing interface (MPI) and can be offloaded to GPUs thanks to dedicated CUDA kernels.

## 1. Motivation and significance

Interactions between proteins that lead to the formation of a three-dimensional (3D) complex is a crucial mechanism that underlies major biological activities ranging from immune defence system to enzyme catalysis. The 3D structure of such complexes provides fundamental insights on the protein recognition mechanism and protein functions [1, 2]. To complement the labor-intensive experimental characterization of protein complexes computational docking approaches have been developed to predict their 3D structures [3, 4]. The prediction of these structures using docking usually consists of two steps: First, the sampling step that consists of systematically (or randomly) rotating and translating individual protein components to generate typically tens of thousands of candidate interaction models; second, the scoring step that evaluates each of the models and selects the ones that are most likely to occur in nature.

The scoring problem has been a highly challenging task for decades. Many methods have been developed and can be largely grouped into five types: 1) Shape complementarity based methods, favoring models that maximize the surface matching with minimal shape penetration [5, 6], 2) physical energy-based methods, which sum up all the pairwise interaction energies between interface atom/residue pairs and are widely used in most modern docking software [7, 8, 9, 10], 3) statistical potential-based methods, which coverts the interaction frequency of interface atom/residue contact pairs observed in the experimentally solved protein complexes to potentials using the Boltzmann distribution [11, 12], 4) machine learning based methods, which typically treat the scoring problem as a binary classification problem, predicting a docked model as near-native or not [13, 14, 15], and 5) co-evolution based methods, which score models based on the co-occurrence frequencies of residue pairs in sequence alignments [16]. Different scoring approaches are regularly benchmarked against each other during a community-wide challenge, the Critical Assessment of Prediction of Interactions (CAPRI) [17].

Recently, we introduced a novel graph kernel based machine-learning approach, called iScore [18]. iScore represents the interface of a protein complex as an interface graph, with the nodes being the interface residues and the edges connecting the residues in contact. By comparing the graph similarity between the query graph and the training graphs, iScore predicts the likelihood how close the query graph is to a near-native model. We have demonstrated in our previous publication [18] that iScore competes with, or even outperforms various state-of-the-art approaches on two independent test sets: the new entries of Docking Benchmark 5.0 set[19] and the CAPRI score set [20]. Using only a small number of features, i.e. 1 evolutionary feature and 3 physics energy terms, iScore performs well compared with IRaPPA[15], the latest machine-learning scoring function, which exploits 91 features. This demonstrates the advantage of representing protein interfaces as graphs as compared to fixed-length feature vectors which discard information about the interaction topology.

We present here the software for iScore. As explained in the following, the software is easy to use thanks to dedicated executable scripts that completely automate the computational workflow. Furthermore, the software leverages distributed and heterogeneous computing technologies to accelerate the generation of the required data and its analysis.

## 2. Software description

The underlying method is described in details in [18] and only a summary is provided here to highlight the different components of the software. As described in [18], the interface of each protein-protein model is represented as a bipartite graph. Each node is labelled with a 20×1 feature vector from the position-specific scoring matrix (PSSM) of the corresponding residue. PSSMs [21] are widely used in bioinformatics and encode the log-likelihood ratio of the observed frequency of each amino acid type at a specific sequence location against a background frequency. They therefore represent the degrees of conservation for the protein’s residues at their specific location in the sequence. The similarity between two graphs is evaluated via a random walk graph kernel (RWGK) approach [22]. The graph-pair similarity matrix is used as input of a support vector machine (SVM) to classify the interface graphs as near-native or non-near native. The decision value of the SVM classification is then combined with energetic terms to score each proteinprotein interface (PPI). As for any supervised learning approach, the SVM model is first trained on a well defined dataset before being used to classify new conformations.

The software presented here provides a fully automatized end-to-end training and testing platform for the ranking of PPIs following the iScore method. The software is organized as a Python module containing dedicated classes in charge of specific steps in the computational workflow. This workflow is fully automatized through executable scripts that orchestrates the entire computation from processing PDB files of the docked models to obtaining the final score of each PPI.

### 2.1. Software Architecture

The general architecture of iScore for training a model and scoring new conformations is represented in Fig. 1. The software only requires PDB files of the docking models contained in the dataset used for training or scoring. All other intermediary files are automatically generated and processed by the software. We provide details in the following different steps of the computational workflow and describe each module.

**Figure 1:**
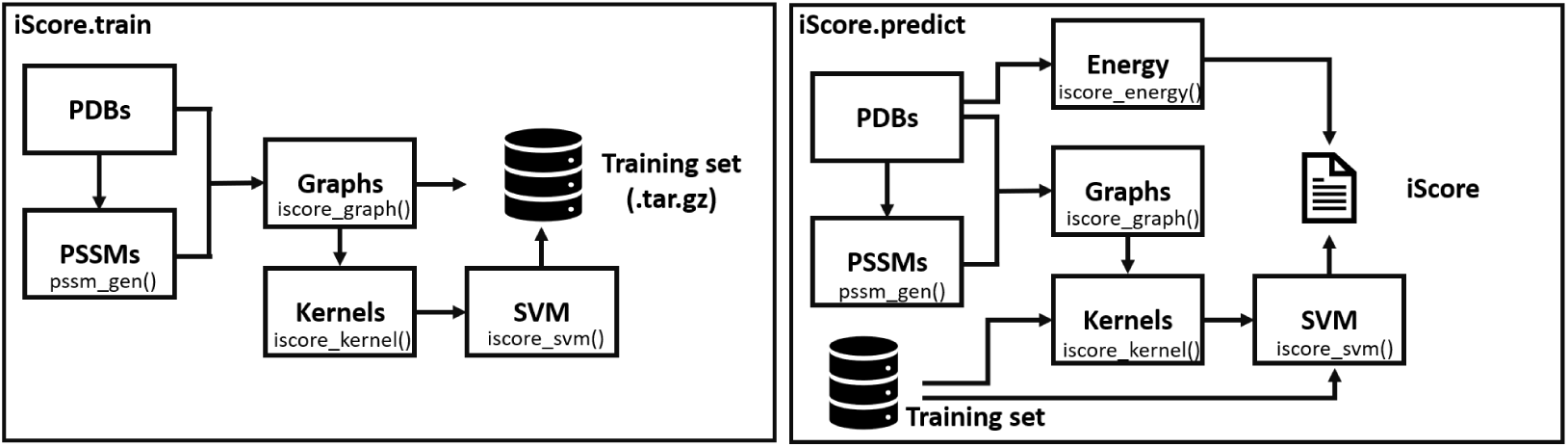
Computational workflow of iScore during the training of a SVM model and during the utilisation of pre-trained model to rank new PPIs.

#### 2.1.1. Generation of the PSSM files

PSSM files of docked conformations are generated by the pssm gen() class using PDB files for input.The calculation of the PSSM relies on PSIBLAST[23] using BLAST version 2.7.1+.The default parameters of the BLAST (for example, substitution matrix, gap costs, etc) were set in agreement with the recommended values provided in the BLAST user guide [24]. Other parameters are provided in [18]. The pssm gen() class also formats the PSSM files for further processing. The class outputs resulting PSSM files for each chain in the PDB files into a separate folder for further processing.

#### 2.1.2. Generation of interface bipartite graphs

The graph generation is handled by the iscore graph() function and relies heavily on our pdb2sql tool that allows manipulating PDB files using SQL queries [25]. The contact residues are identified by the interface module of pdb2sql using a default contact distance of 6.0 Å. The PSSM files generated in the previous step are then read and checked against the sequence of the protein. The PSSMs are subsequently mapped onto the interface graph. The resulting graph is then serialized using the pickle library in order to exploit the object hierarchy in the next computational steps. The class also provides options to export multiple graphs in a single HDF5 file for further visual inspection (see section 3).

#### 2.1.3. Random walk graph kernels

The function iscore kernel() is responsible for the computation of pairwise random-walk graph kernels. For each pair of PPIs contained in the dataset, the corresponding graph files are first unpickled and loaded in memory. The different elements necessary to the computation of the RWGK are computed and assembled in the final kernel value (see [18] for details on the calculation). All kernel values are then stored in a dedicated pickle file.

#### 2.1.4. Training the SVM model

The function iscore svm() can then be used to train an SVM model from the previously computed RWGK. To this end, users must also provide the ground truth, i.e. the binary class 0/1 of each conformation contained in the training dataset. In iScore, we choose the binary label 1(0) to describe a near-native(non near-native) conformations. The function relies on the libSVM library [26] to train the SVM model. To facilitate the further exploitation of the trained SVM model, the SVM model are efficiently packed into a dedicated archive together with the graphs of all the conformations of the training set. This self contained archive contains all the information required to score new PPIs.

#### 2.1.5. Scoring new PPIs

The workflow for ranking new PPIs is very similar to the one used to train the SVM model. Users only need to provide PDB files of new conformations and compute the corresponding PSSMs and interface graphs. However, the RWGK are now computed between the new conformations and the ones contained in the training set. This can easily be done using the training archive that contains all the relevant information. The resulting kernels are then used as input for the SVM model, and the SVM decision value is used as one component of the final scoring function. The other component of the scoring function is provided from energy terms that are directly computed from the PDB files of the new conformations. The weight of each term in the final scoring function has been optimized using genetic algorithm as explained in [18].

### 2.2. Software functionalities

Beyond the individual modules described above, the software provides crucial functions that facilitate and accelerate the process of ranking PPIs.

#### 2.2.1. Automation of the Computational Workflows

The software provides executable scripts that fully automatize the workflows illustrated in Fig. 1. These scripts seamlessly orchestrate all the computational steps at the exception of the calculation of the PSSMs. This calculation can be rather demanding and therefore must be performed as a pre-processing step using the functions provided by the software.

The training an SVM model from the PDB leads to the creation of a training archive and is fully controlled by the iScore.train.mpi executable script. This script reads the PDB and PSSM files, generates the interface graphs, computes all the pairwise RWGKs, trains the SVM model, and finally assembles the training archive. Similarly, the ranking of new conformations using a trained model can simply be achieved via a single command: iS-core.predict.mpi. This script reads the PDB and PSSM files, generates all the interface graphs, computes the RWGK between the new conformations and the conformations included in the training set, and finally scores the new conformations.

To face the potentially large computational cost associated with the calculation of the interface graphs and their pairwise RWGKs, these executable scripts support the distribution of the computational load across different MPI processes using mpi4py [27]. For the calculation of the graphs, the different conformations are distributed among the different MPI processes, and for the RWGK calculations all the pairwise combinations are distributed among the MPI processes. Simple performance benchmarks are reported in Fig. 2a showing good performance of the MPI distribution. However, note that training the SVM model and scoring new PPIs are done using a single process.

**Figure 2:**
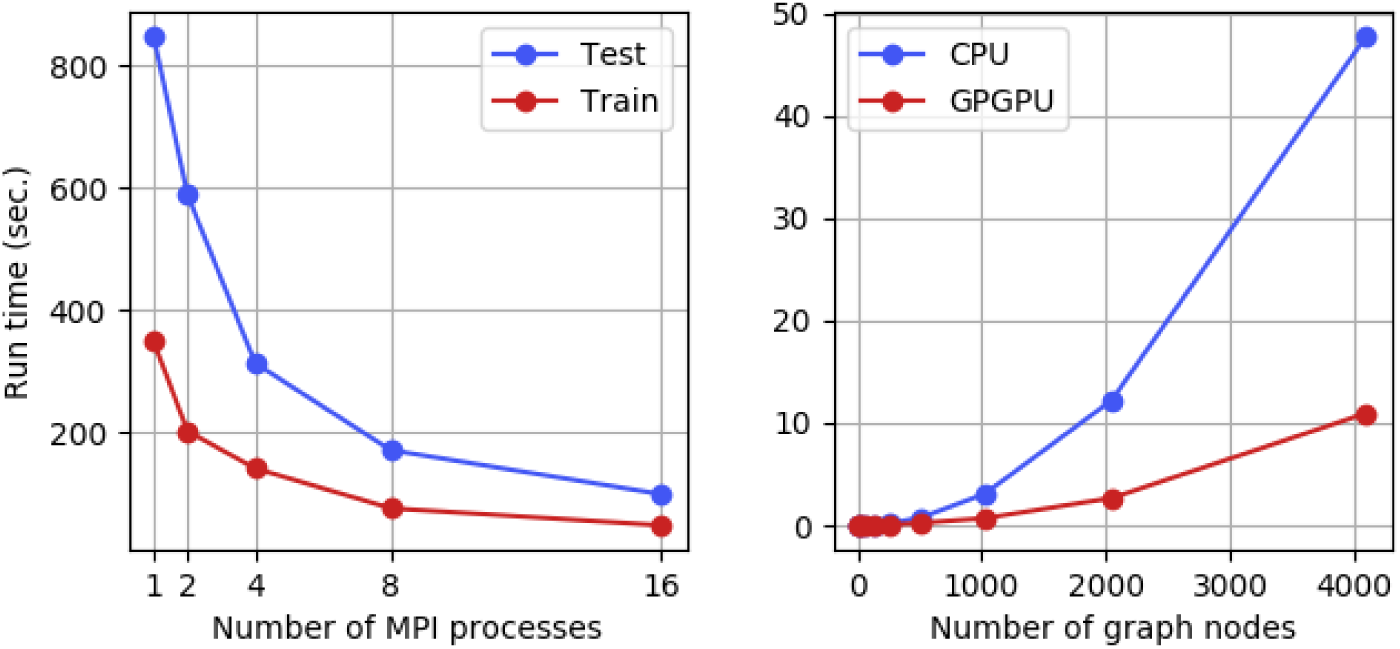
a) Scaling of iScore.train.mpi and iScore.predict.mpi with respect to the number of MPI processes. The training and testing set contained 234 and 599 conformations respectively. b) Average run time on CPU (Intel Xeon E5-2650 v4 @ 2.20 GHz) and GPU (Nvidia GeForce GTX 1080 Ti) for the calculation of RWGK for two graphs containing *n* nodes and 3*n* edges.

#### 2.2.2. Calculation of the RWKG on GPUs

To accelerate the calculations of the graph kernels, we have developed simple GPGPU kernels using pyCUDA [28]. The utilization of these GPGPU routines can easily be turned ON or OFF through one optional keyword argument of the iscore kernel() function. Fig. 2b shows the runtime of the CPU and GPU routines computing the RWGKs. As seen on this figure, a sizable improvement of performance can be obtained for large graphs. The software also provides solutions to tune the GPU kernels through the kernel tuner library [29]. This allows to automatically find the optimal configuration of the kernel in terms of blocs size, threads size, etc.

While the GPU routines might be interesting to process very large proteins or for other applications, we have exclusively used the CPU routines in our calculations because our protein interface graphs contain less than a hundred nodes.

#### 2.2.3. Visualization

As mentioned in section 2.1.2, the interface graphs computed by iScore can be stored in a HDF5 file for further analysis. The resulting HDF5 file contains an entry for each graph where all the relevant data are stored. To facilitate the inspection and exploration of these interface graphs, we have developed a simple graphical interface based on the customizable HDF5 browser h5X [30]. This interface is accessible via the executable iScore.h5x. This interface allows to quickly generate all the data for visualization of a given graph connection using PyMol [31]. An example of representation is shown in Fig. 3. This figure shows a single PPI. All the contact residues are highlighted by a stick representation and bright color whereas the rest of the protein structure is represented by thin grey lines. Edges linking the contact residues represent the contact between the two chains and the label of each contact residue is displayed for clarity.

**Figure 3:**
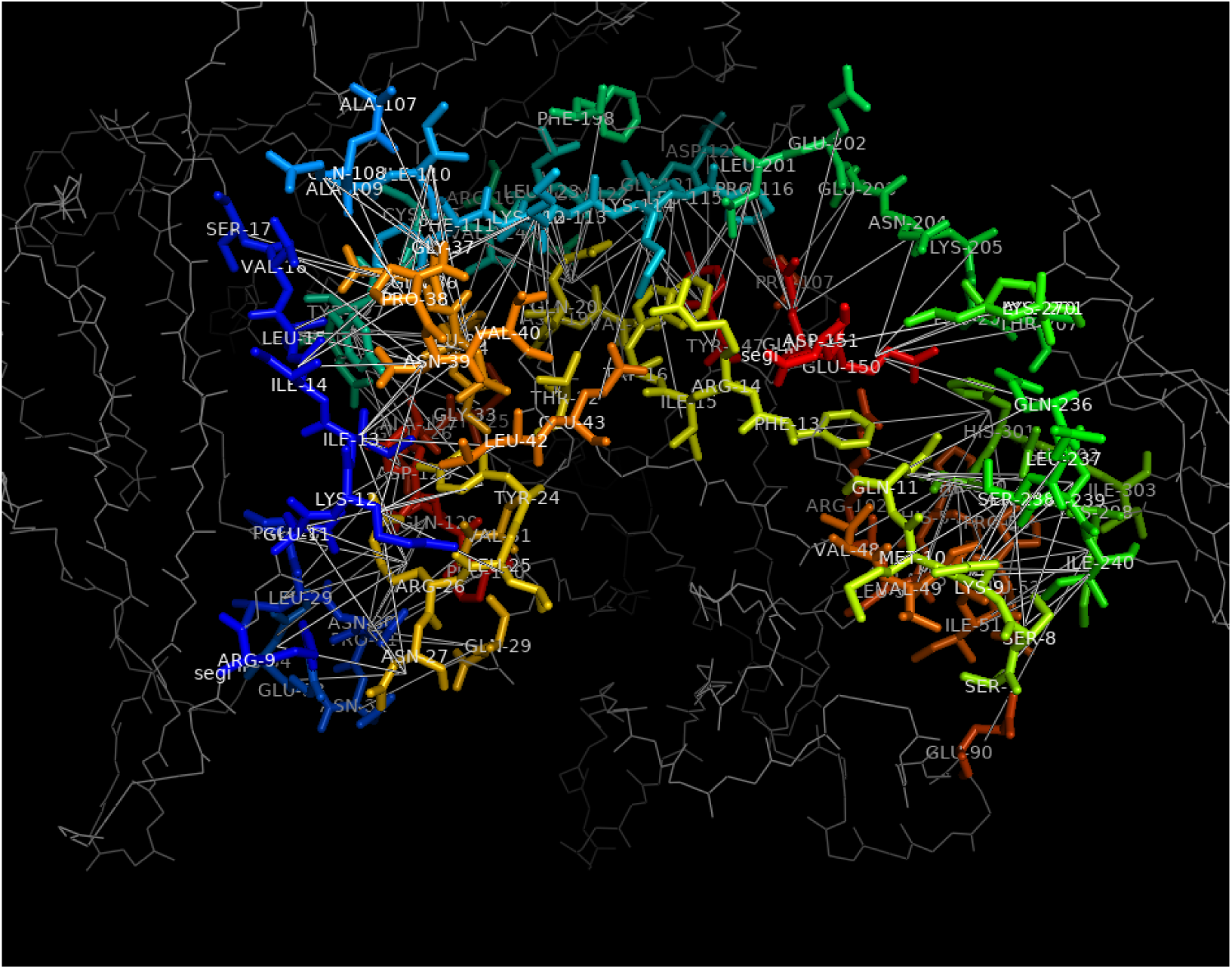
Visualisation of the connection graph of PDB ID: 1IRA using iScore.h5x with the PyMol molecular viewer.

### 2.3. Code Snippets

Beyond the executable scripts mentioned above, iScore can also be used as a Python module and could therefore be integrated in other applications.

We illustrate here the use of iScore through a small code snippet.

~~~
1 from iScore. graphrank. graph import GenGraph
2 from iScore. graphrank. kernel import Kernel
3
4 # generate the first graph
5 pdb = ‘ 1ATN .pdb ‘
6 pssm = {‘A‘:‘ 1ATN.A.pdb. pssm ‘,
7         ‘B‘:‘1ATN .B.pdb. pssm ‘}
8 gen = GenGraph (pdb,  pssm)
9 G1 = gen. get_graph ()
10
11 # generate the second graph
12 pdb = ‘1IRA .pdb ‘
13 pssm = {‘A‘:‘1IRA .A.pdb. pssm ‘,
14         ‘B‘:‘1IRA .B.pdb. pssm ‘}
15 gen = GenGraph (pdb,  pssm)
16 G2 = gen. get_graph ()
17
18 # compute the kernel
19 K = Kernel ()
20 K. compute_kron_mat (G1, G2)
21 K. compute_px (G1, G2)
22 K. compute_W0 (G1, G2)
23 ker = K. compute_K (lamb =1.0,  walk =4)
~~~

As we can see on this snippet, iScore provides the solution to generate graphs of given structures using PSSM as a node attribute and to compute the random walk graph kernel between the graphs. The graph generation is done via the GenGraph() class that only takes PDB and PSSM files as input. The random walk graph kernel of the two graphs can then be computed using the Kernel() class and its methods.

## 3. Illustrative Examples

We present here the results on test cases extracted from previous CAPRI competitions. In order to score the conformations contained in the test cases, a representative training set containing 234 distinct PPIs was first assembled. Half of these conformations correspond to real experimental structures of complexes chosen from the Docking Benchmark 4 (DB4) [32]. The second half correspond to non-native docking models generated using the HAD-DOCK docking software from entries of the DB4. Their i-RMSD values are larger than 10 Å. The resulting dataset is publicly available [33].

The trained model was used to score and rank conformations from previous CAPRI targets, namely targets T32, T41, T47 and T50. Fig. 4 shows the corresponding hit rate plot obtained with iScore and the HADDOCK scoring function. Hit rate plots are commonly used to compare different scoring functions. The hit rate at *N* represents the fraction of near-native models contained in the best *N* models predicted by a scoring function. As seen in Figure 4 and Table 1, iScore performs better than HADDOCK on 2 of these cases (T32 and T41) and shows similar performance on the remaining two. These results are in line with those reported in [18] where iScore performed very well on a large range of test cases.

**Table 1:**
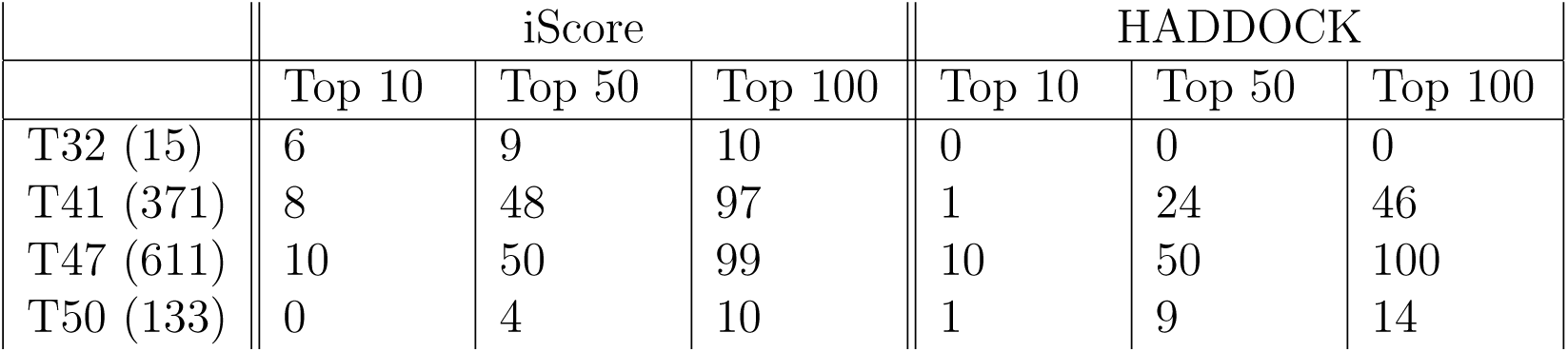
Performance of iScore and HADDOCK scoring functions on four CAPRI test cases. The number in bracket represents the number of near-native conformations for each case. The number in each column represents the number of near-native conformations in the top 10, top 50 and top 100 conformations predicted by the two methods.

**Figure 4:**
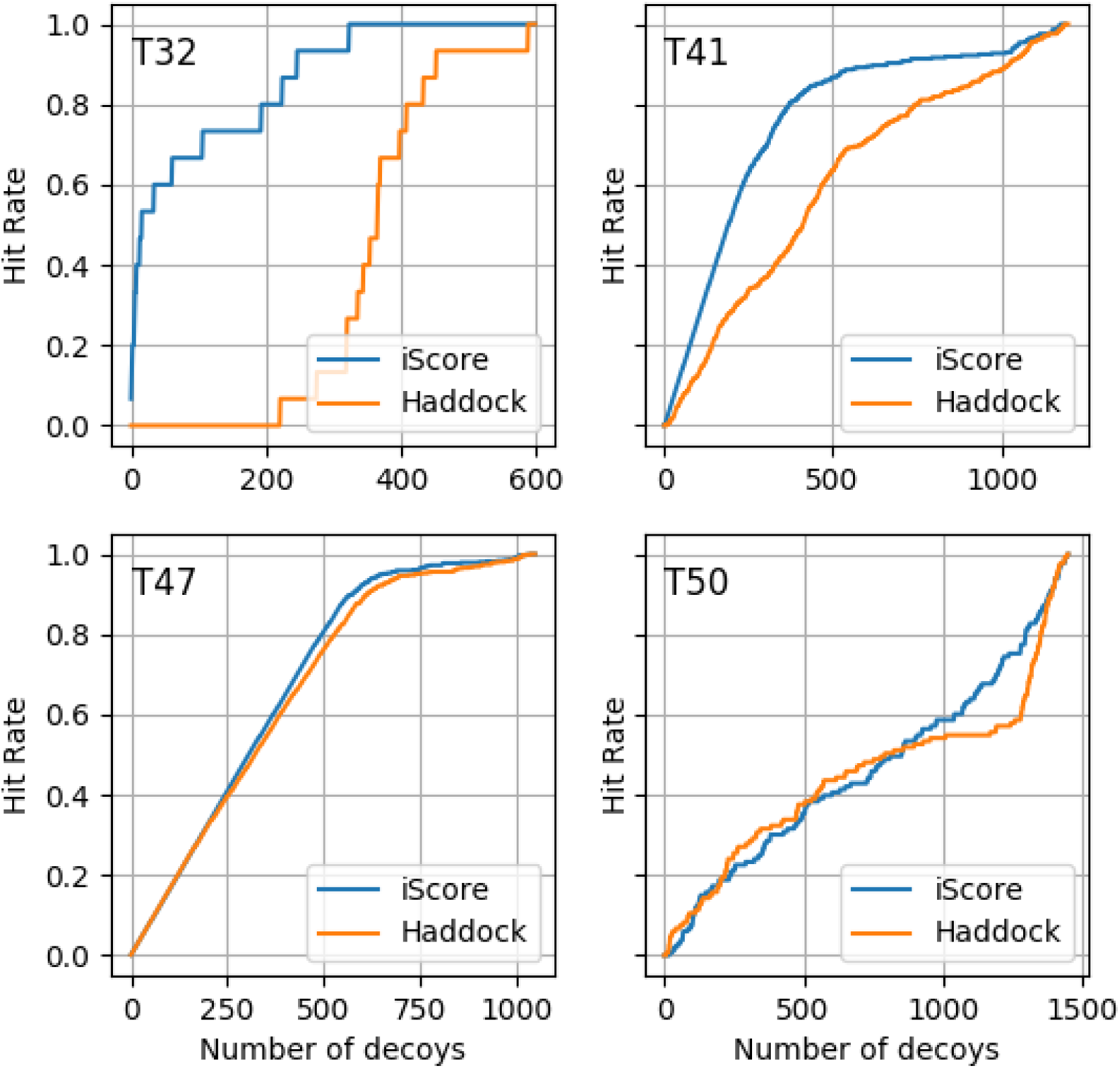
Comparison of the hit rates obtained by iScore and HADDOCK for four CAPRI targets.

## 4. Impact

The software presented in this paper provides ease of use in end-to-end platform for scoring and ranking of PPIs. Thanks to the provided executable scripts, users can easily generate the graphs, compute their pairwise kernels and use them to train a SVM model. The self-contained archive file generated during the training contains all the necessary information to rank new docking conformations. This enables to simplify data handling and facilitates the exchange of trained model between different users. The dedicated scripts briefly described in section 2.2.1 fully automatize the computational workflows supporting the training and testing of a SVM a model. This workflow not only makes the use of the code easier, therefore facilitating its adoption by the community, but also ensures more reproducibility. The clear architecture of the software facilitates its maintenance and further development.

The distribution of the computational load using MPI significantly reduces the time for training and using SVM models: Training our SVM model used in section 3 takes under 50 seconds using 16 cores while scoring the 600 conformations contained in the T32 CAPRI test case takes less than 2 minutes.

The software presented in this paper has already been used in a recently published paper that describes the underlying methodology and used it on a large range of test cases. In agreement with Fig. 4, the results presented in [18] are competitive compared to widely used scoring functions such as HADDOCK. The software also recently has been used during the CAPRI competition.

While the software has been developed specifically for ranking PPIs, the method is generic and may be used for any problem where graphs have to be ranked. To this end, the software could be generalized for users to specify the nodes and edges features as well as the kernel used in the definition of the graph kernels. This generalization will be done in a near future.

## 5. Conclusions

We have presented a new software package, iScore, that provides an end-to-end solution for ranking protein-protein interfaces. The method is based on a support vector machine classification using random-walk graph kernels as input. The software is built as a Python package that can be used either interactively or through the use of dedicated executable scripts that fully automatize the computational workflow. The calculations can be distributed across multiple MPI processes and GPGPU kernels have been developed to accelerate the calculation of graph kernels. As a whole, the software provides a user friendly solution for ranking PPIs more efficiently and accurately.

## Acknowledgements

This work was supported by an Accelerating Scientific Discovery (ASDI) grant from the Netherlands eScience Center (grant no. 027016G04). CG acknowledges financial support from the China Scholarship Council (grant no. 201406220132). LX acknowledges financial support from the Netherlands Organisation for Scientific Research (Veni grant 722.014.005) and from an Accelerating Scientific Discovery (ASDI) grant from the Netherlands eScience Center (grant no. 027016G04). VH acknowledges financial support from the National Science Foundation (grant no. ACI 1640834) and the National Institutes of Health (NCATS UL1 TR002014-01), the Center for Big Data Analytics and Discovery Informatics which is co-sponsored by the Institute for Cyberscience, the Huck Institutes of the Life Sciences, and the Social Science Research Institute at the Pennsylvania State University, and the Edward Frymoyer Endowed Professorship at Pennsylvania State University and the Sudha Murty Distinguished Visiting Chair in Neurocomputing and Data Science sponsored by the Pratiksha Trust at the Indian Institute of Science.

## Required Metadata

### Current code version

**Table 2:** Code metadata

| Nr. | Code metadata description | Please fill in this column |
| --- | --- | --- |
| C1 | Current code version | 0.2.0 |
| C2 | Permanent link to code/repository used for this code version | https://github.com/DeepRank/iScore |
| C3 | Legal Code License | Apache-2.0 |
| C4 | Code versioning system used | git |
| C5 | Software code languages, tools, and services used | python, MPI, CUDA. |
| C6 | Compilation requirements, operating environments & dependencies | numpy, libSVM, pdb2sql, h5x, pytest, biopython, mpi4py, numpy, scipy, h5py, matplotlib |
| C7 | If available Link to developer documentation/manual | https://iscoredoc.readthedocs.io/ |
| C8 | Support email for questions |  |

